# Willingness to pay for spectacle among presbyopic population in South Gondar, Ethiopia: A cross-sectional study

**DOI:** 10.1101/482943

**Authors:** Haile Woretaw Alemu

## Abstract

**Aim:** To assess the willingness to pay for spectacles among the south Gondar presbyopic adult population.

**Methods:** An interview-based questionnaire was employed to elicit the willingness to pay for spectacles among people refracted during an outreach service at Debre-Tabor Zonal Hospital.

**Results:** Of the total 322 people participating in the study, only 53.4% (172) were experienced spectacles users. The median gross monthly income of participants was US$ 75.0 (ranged US$ 7.1 - 321.4) and the mean amount of money willing to pa for a pair of spectacles was US$ 17.9 (ranged US$ 1.1-107.1). Participants who were willing to pay US$ 12.5 for a pair of spectacles from a government optical accounted 63.0% (95% CI: 57.8-68.3), while those willing to pay the minimum international pair of spectacle price US$ 5.6 were accounted 73.9% (95% CI: 68.9-79.2%) and spectacle from local private optical price US$ 17.8 accounted 46.6% (95% CI: 40.4-52.2). Multivariate logistic regression analysis indicated factors such as age (P=0.049), occupation (0.001), monthly income (0.001) and history of previous spectacle wear (0.005) to be significantly associated with willingness to pay for a pair of spectacles.

**Conclusion:** Determining willingness to pay and affordable price could increase spectacle coverage among presbyopic individuals. confirmatory public willingness to pay for a pair of spectacles has to be considered as a useful aspect in establishing affordable and a financially sustainable optical service. The positive willingness to pay for a pair of spectacle has to be supported with accessible provision of spectacles.

## Introduction

Presbyopia is a physiological, age-related change leads to loss of lens accommodation and an inability to focus at near.^1^ Commonly presbyopia manifest between the ages of 40-45 years^2-4^ but still reports confirm occurrence as early as 30 years of age.^1,5-7^

An estimated 1.272 billion people are presbyopic in the world, of which 244 million are uncorrected and reach up to 94% of cases in lower-income countries. It affects working age group and contributes an estimated 25.367 billion dollars productivity loss.^8,9^ In sub-Saharan Africa, the proportion of visual impairment due to uncorrected refractive error for presbyopic age group ranged from 12.3% to 57.1%.^10^ A population-based studies in East Africa reported the prevalence of presbyopia range from 61.7%–85.4%.^11,12^ Similarly, the prevalence rate of presbyopia was 62% in rural Tanzania^13^ and 68.7% in Ethiopia.^14^ Therefore, it clearly poses an important public health challenge.^1^

The mainstay of presbyopia correction is spectacle. It is an effective, economic option for low and middle-income countries. In Rwanda 95% cases could manage with spectacle with cost range from $ 0.46 to 4 US dollar hence with provision of spectacle correction predicted average productivity gains were estimated 10% of GDP.^15^ According to Patel et al. a high proportion of people (69%) especially men were able to afford spectacles at a price that covered including shipping cost.^1^ Similarly, the mean cost of spectacle in different states of Zambia ranges from US$ 4.0-20.0.^16^

Few population-based studies have been conducted to estimate the willingness to pay for refractive services and spectacles using different methods such as open-ended questions^17^ or binary-with-follow-up.^18^ For spectacle about 82.9% study participants were willing to pay at least US$ 0.10 in Timor-Leste^19^ and 76.6% were willing to pay at least US$ 0.38 in Cambodia.^18^ Likewise, in Zanzibar the mean amount willing to pay for spectacles was US$ 3.14.^20^

Centralized and limited spectacle manufacturing workshop, additional costs (such as transport) and undetermined willingness to pay for a pair of spectacles challenges presbyopia intervention program. Therefore, this study assessed willingness to pay for a pair of spectacles among presbyopic population of the South Gondar administrative zone.

## Methods and Materials

A cross sectional study design was implemented. This study was conducted between January to march 2018 in the South Gondar zone. Study area has a single eye care centre without spectacle workshop service. Spectacles provision is through visiting outreach service in partnership with the University of Gondar and Vision Aid Overseas.

During outreach service a total of 3220 subjects screened and of this 1018 were presbyopic. Subjects who have presbyopia, underwent for complete eye examination and willing to use spectacle were included in the study. the determined sample size was 340 subjects and selected using simple random sampling technique using single proportion formula by considering 82.9% proportion and 4% marginal error.

Pre-tests were taken to refine and validate the questionnaire. Using structured questioner optometrists collected information such as demographic and socioeconomic data, history of spectacles uses and future source preferences. Willingness to pay was determined by comparing participants stated willingness to pay against minimum local government opticals spectacle price US $12.5. Each study participant undergone vision screening, complete eye examination and refractive assessment by optometrists. Presenting unaided distance and near visions was taken in a suitably lit room for each eye separately using Snellen 6-meter chart for distance and reduced Snellen reading E-chart for near vision. Presenting distance vision of less than 6/18 in the better eye was considered as visual impairment. Presbyopia was defined (when wearing a distance correction if needed) as a presenting near vision acuity worse than N8 at 40 centimeters or if an addition of at least +1.00DS was required to improve near vision to at least N8.

Data was entered and analyzed using Statistical Package for Social Science (SPSS version 20, Chicago, IL, USA). Descriptive statistics were summarized with the same package. Significance was set at less than 5%. willingness to pay for spectacles was analyzed using logistic regression to determine associated factors. One-way ANOVA and one-sample t-test were used to compare mean difference of willingness to pay across sources. The tenets of the declaration of Helsinki were observed and ethical approval was obtained from College of Medicine and Health Science ethical review committee, University of Gondar. Informed consent was obtained from each participant prior to questionnaire administration.

## Results

Out of 340 recruited subjects 322 gave valid and complete response and included in the analysis. The mean age of study participants was 51 ± 7.8 years (ranged 35 to 73). Of the total 322 subjects, 73.3% (236) were male, 72.4% (233) were urban residents and 46.0% (148) were government employees.

In addition to presbyopia, 14.6% (47) subjects were visually impaired at distance. Subjects who were aware of the existence of eye care services in the town accounted for 84.8% (273) but only 58.4% (188) had a history of previous eye examination.

In terms of previous history of using spectacle, only 53.4% (172) of the study participants were experienced users. Within that group, those who obtained spectacles from street sellers accounted for 38.4% (66), outreach suppliers 25.0% (43), private optical outlets 25.0% (43) and government optical outlets 11.6% (20). Whereas, in the future, if spectacle wear was recommended, 51.9% (167) subjects would prefer to use government owned optical outlets, followed by 45.0% (145) from outreach services and 3.1% (10) from private opticals. The motives behind these choices were to seek better quality in 49.1% (158) followed by cost in 47.2% (152) and fast supply in 3.7% (12). Table 1: sociodemographic and economic characteristics of study participants

**Table 1:.** sociodemographic and economic characteristics of study part cipants

| *Variables* |  | *Frequency (N=322)* | *Percent (%)* |
| --- | --- | --- | --- |
| Gender | Male | 236 | 73.3 |
|  | Female | 86 | 26.7 |
| Age category (years) | ≤49 | 138 | 42.9 |
|  | 50-59 | 130 | 40.4 |
|  | ≥60 | 54 | 16.8 |
| Residential area | Urban | 233 | 72.4 |
|  | Rural | 89 | 27.6 |
| Religion | Christian | 294 | 91.3 |
|  | Muslim | 28 | 8.7 |
| Educational status | Literate | 293 | 91.0 |
|  | Illiterate | 29 | 9.0 |
| Occupation | Gove employ | 148 | 46.0 |
|  | Private business | 59 | 18.9 |
|  | Farmer | 62 | 18.6 |
|  | *Others | 53 | 16.5 |
| Household monthly Income (US\$) | ≤53.6 | 112 | 34.8 |
|  | 53.61-107.1 | 127 | 39.4 |
|  | ≥107.2 | 83 | 25.8 |
| Presence of visual impairment | Yes | 47 | 14.6 |
|  | No | 275 | 85.4 |
| Previous eye examination | Yes | 188 | 58.4 |
|  | No | 134 | 41.6 |
| Aware of eye care service availability | Yes | 273 | 84.8 |
|  | No | 49 | 15.2 |
| Previous spectacle wears | Yes | 172 | 53.4 |
|  | No | 150 | 46.6 |

\*Others (Daily laborer, Housewife, Retired)

The median gross monthly income of participants was US$ 75.0 (range US$ 7.1 - 321.4). Of the total study participants 63.0% (95% CI, 57.8-68.3) were willing to pay a minimum local government optical spectacle price of US$ 12.5. While 73.9% (95%CI, 68.9-79.2%) participants could afford a minimum international pair of spectacles at US$ 5.6, only 46.6% (95% CI, 40.4-52.2) could afford a local private spectacle priced at US$ 17.8. The mean amount of money the study participants were willing to pay for a pair of spectacles was US$17.9 (range US$1.1-107.1). The difference between this mean and that of the international spectacle price of US$ 5.6 and local government optical spectacle price $12.5 USD was US$ 12.25 (p=0.003, 95% CI, 10.8-14.0) and US$ 5.4 (p=0.003, 95% CI, 3.8-7.2) respectively.

The ANOVA test showed that the mean amount of money willing to pay for a pair of spectacles among optical sources varies significantly (p =0.0001). The mean difference of willingness to pay for outreach supplies compared to the government optical outlets was US$ 12.5 (p=0.0001, 95% CI, 8.9-16.2) and the mean difference in outreach supply compared with private optical outlets was US$ 18.1 (p=0.0001. 95% CI, 7.6-28.6).

Multivariate logistic regression analysis indicated factors such as gender, religion, place of residence, visual status, history of eye examination and awareness of eye care services availability had no association with willingness to pay for spectacle. While factors such as age (P=0.049), occupation (0.001), monthly income (0.001) and history of previous spectacle wear (0.005) was significantly associated with willingness to pay.

Participants within the age group ≤49 years were 6.2 times (CI:1.526-11.353) and age group from 50-59 years were 5.9 times (CI: 1.341-25.907) more willing to pay compared to those in the age group of 60 years and above. With respect to occupation, government employees were 7.2 times more willing to pay (CI: 1.748-30.042) than the rest of the cohort.

Household monthly income was significantly associated with willingness to pay for spectacles. Hence participants who earned between US$ 53.6-107.0 were 24 times (CI: 8.600-66.272) and those who earned more than US$ 107.1 were 48 times (CI: 8.591-71.758) more willing to pay compared to those who earned less than US$ 53.0.

Participants who were experienced spectacle users were 4 times (CI: 1.526-11.353) more willing to pay when compared to neophytes. Table 2: Associated factors with willingness to pay the minimum local government spectacle price.

**Table 2:** Associated factors with willingness to pay the minimum local government spectacle price.

| Variable | Willingness to pay US\$ 12.5 | | COR | AOR |
| --- | --- | --- | --- | --- |
|  | Willing | Unwilling |  |  |
| **Gender: Male** | 152(64.4%) | 84 (35.6%) | 1.242(0.749-2.060) | 2.194(.787-6.116) |
| **Female** | 51(59.3%) | 35 (40.7%) | 1 | 1 |
| **Age(years)** |  |  |  | 0.049* |
| **≤49** | 89 (64.5%) | 49 (35.5%) | 2.448(1.288-4.653) | 6.287 (1.231-32.110)* |
| **50-59** | 91 (70.0%) | 39 (30.0%) | 3.145 (1.630-6.067) | 5.894 (1.341-25.907)* |
| **≥60** | 23 (42.6%) | 31 (57.4%) | 1 | 1 |
| **Residency** |  |  |  |  |
| **Urban** | 184 (79.0%) | 49 (21.0%) | 0.072(0.040-0.131) | 4.337(0.912-20.627) |
| **Rural** | 19 (21.3%) | 70 (78.7%) | 1 | 1 |
| **Religion** |  |  |  |  |
| **Christian** | 196 (66.7%) | 98 (33.3%) | 6.00 (2.466-14.598) | 2.862 (0.528-15.503) |
| **Muslim** | 7 (25.0%) | 21 (75.0%) | 1 | 1 |
| **Occupation** |  |  |  | 0.01* |
| **Gov employ** | 140 (94.6%) | 8 (5.4%) | 53.846 (20.861-138.985) | 7.247 (1.748-30.042)* |
| **Private B** | 41 (67.2%) | 18 (32.8%) | 6.308 (2.770-14.364) | 1.880 (0.476-7.418) |
| **Farmer** | 9 (15.0%) | 53 (85.0%) | 0.543(0.211-1.397) | 0.605 (0.081-4.508) |
| **Others** | 13 (24.5%) | 40 (75.5%) | 1 | 1 |
| **HM income** |  |  |  | 0.0001* |
| **≤1500** | 18 (16.1%) | 94 (83.9%) | 1 | 1 |
| **1501-3000** | 104 (81.9%) | 23 (18.1%) | 23.614(12.000-46.467) | 23.873 (8.600-66.272)* |
| **≥3001** | 81 (97.6%) | 2 (2.4%) | 211.500(47.629-939.181) | 48.319 (8.591-71.758)* |
| **Presence of VI** |  |  |  |  |
| **Yes** | 17 (36.2%) | 30 (63.8%) | 0.271 (0.142-0.518) | 0.932 (0.270-3.218) |
| **No** | 186 (67.6%) | 89 (32.4%) | 1 | 1 |

## Discussion

Although different method makes comparison difficult, in this study spectacle coverage was twice as high as in a Zanzibar study.^20^ Inclusion of more urban resident and literate study participants could increase the coverage. Spectacle wear was higher among literate, male, government employees, urban residents and age group from 50-59 years. Spectacles were mainly obtained from unauthorized street vendors. Although they are easily available they are of poor quality. There is limited access either to government or privately owned optical outlets. However, in future, study participants showed preference to get their spectacles from government owned optical outlets anticipating quality spectacle with fair costs.

The mean amount of money participants was willing to pay for a pair of spectacles was US$ 19.0 (ranged from US$ 1.1 to111.0). It was nearly 3 times higher than the standard international cost of spectacles (US $ 5.7 equivalent with US$ 3.0 in 2006) and 1.5 times higher than local government optical outlet’s price. However, there was no neither government or nor privately-owned optical outlets available in the study area. Hence it costs an extra $ 10.0 USD for transport to get access to optical shops 100 kilometers away neighboring towns with a week-long waiting period. This constraint could be a major reason why communities have to depend on outreach spectacle supply and street vendors.

More than two thirds of study participants were willing to pay a quarter of their monthly income for a pair of spectacles. In this study, the proportion of willingness to pay was above that found in a study done in Timor-Leste (31.6%).^19^ Similarly, the mean paying capacity was above that found in other studies.^18-20^ This could be explained by participants different socio-economic status, Literacy and variety implementation methods. However, the cost of a pair of spectacles at a government owned optical outlet was higher than the price quoted in the studies of Heidi RL et al ^20^ and Jacqueline et al.^19^ This might be because of limited availability options, extended custom procedures for importation and the lack of optical production sites.

Being in the age group of less than 60 years of age and being a government employee were independently associated with a willingness to pay for a pair of spectacles. This correlates well with studies conducted in Cambodia^18^ and Timor-Leste.^19^ Higher salaries, higher near visual demand and an independent income source are likely to increases the willingness to pay. A monthly income greater than $ 53.5 USD is independently associated with a willingness to pay for a pair of spectacles. This finding is supported by previous studies.^18,19^ Experienced spectacle users were independently associated with a willingness to pay for a pair of spectacles which might be because of a better understanding of the benefits of using spectacles for close tasks. However, this was contradicted by the Cambodia^18^ and Timor-Leste^19^ studies.

The opportunity afforded by participants’ willingness to pay for a pair of spectacles should be utilized to establish sustainable optical service in order to increase presbyopic spectacle coverage with quality and affordable spectacles. Providing refraction service and spectacle prescriptions alone cannot reverse poor spectacle coverage and assure quality unless optical workshop services are compulsorily linked with refractive services.

## Limitations

Although attempts were made to limit responder bias during the interviews, questions may not guarantee validated responses to assess willingness and exact amount to pay. Being conducted on an outreach site might inflated spectacle coverage and is likely to increase the proportion of willingness to pay.

## Acknowledgment

I would like to thank Optometry department, University of Gondar for material support. I also want to extend my gratitude to Mr. Mohammed Seid for his assistance during statistical analysis and Mrs. Margaret Laurence for her language editorial support.

## References

1. Patel I, West SK. Presbyopia: Prevalence, Impact and Intervention. J. Comm. Eye Health. 2007; 20:40–41

2. Weale RA. Epidemiology of refractive errors and presbyopia. Surv Ophthalmol. 2003; 48:515–43.

3. Pandelis A, Alexandros P. Current Management of Presbyopia. Middle East Afr J Ophthalmol. 2014; 21:10–17.

4. A. Hickenbotham, A. Roorda, C. Steinmaus, A. Glasser. Meta-Analysis of Sex Differences in Presbyopia. Investigative Ophthalmology & Visual Science. 2012; 53: 3215–20.

5. Nirmalan PK, Krishnaiah S, Shamanna BR, Rao GN, Thomas R. A population-based assessment of presbyopia in the state of Andhra Pradesh, south India: the Andhra Pradesh Eye Disease Study. Invest Ophthalmol Vis Sci. 2006;47: 2324–8.

6. Uche JN, Ezegwui IR, Uche E, Onwasigwe EN, Umeh RE, Onwasigwe CN. Prevalence of presbyopia in a rural African community. Rural Remote Health. 2014;14:2731.

7. Ajibode HA, Fakolujo VO, Onabolu OO, Jagun O, Ogunlesi TA, Abiodun OA. A community-based prevalence of presbyopia and spectacle coverage in southwest Nigeria. J West Afr Coll Surg. 2016; 6: 66–82.

8. Frick KD, Joy SM, Wilson DA, Naidoo KS, Holden BA. The global burden of potential productivity loss from uncorrected presbyopia. Ophthalmology. 2015;122: 1706–1710.

9. Holden BA, Fricke TR, Ho SM, Wong R, Schlenther G, Cronjé S, et al. Global vision impairment due to uncorrected presbyopia. Arch Ophthalmol. 2008; 126:1731–9

10. Sherwin JC, Lewallen S, Courtright P. Blindness and visual impairment due to uncorrected refractive error in sub-Saharan Africa: review of recent population-based studies. Br J Ophthalmol. 2012 Jul;96(7):927–30.

11. Burke AG, Patel I, Beatriz M, et al. Population-based study of presbyopia in rural Tanzania. Ophthalmology. 2006; 113:723–727.

12. Sherwin JC, Keeffe JE, Kuper H, Islam FM, Muller A, Mathenge W. Functional presbyopia in a rural Kenyan population: the unmet presbyopic need. Clin Experiment Ophthalmol. 2008; 36:245–251.

13. Patel I, Munoz B, Burke AG, Kayongoya A, McHiwa W, Schwarzwalder AW, West SK. Impact of presbyopia on quality of life in a rural African setting. Ophthalmology. 2006;113(5):728–34.

14. Andualem HB, Assefa NL, Weldemichael DZ, Tefera TK. Prevalence and associated factors of presbyopia among school teachers in Gondar city, Northwest Ethiopia. Clin Optom (Auckl). 2017; 20,9:85–90

15. Vision for a Nation Foundation. Primary Eye Care Service in Rwanda: Benefits and Costs. Croock Associates September 2015. https://d2fyyic8pcxcmc.cloudfront.net/documents/127-3828-vfan-cba-iii.pdf

16. K Griffiths, M Bozzani, A Gheorghe, L Mwenge, C Gilbert. Cost effectiveness of eye care services in Zambia. Cost Eff Resour Alloc. 2014; 12: 6.

17. Onwujekwe O, Uzochukwu B. Stated and actual altruistic willingness to pay for insecticide-treated nets in Nigeria: validity of open-ended and binary with follow-up questions. Health Econ 2004; 13:477–92.

18. J Ramke, A Palagyi, R du Toit, G Brian. Using assessment of willingness to pay to improve a Cambodian spectacle service. Br J Ophthalmol. 2008; 92:170–174.

19. Jacqueline R, Anna P, Rénée dT, Garry B. Stated and Actual Willingness to Pay for Spectacles in Timor-Leste. Ophthalmic Epidemiology. 2009; 16:4, 224–230

20. Heidi RL, Fatma O, Hakika J, Garnia K, Clare G. Presbyopic Spectacle Coverage, Willingness to Pay for Near Correction, and the Impact of Correcting Uncorrected Presbyopia in Adults in Zanzibar, East Africa. IOVS. 2010; 51:2,1234–41

